# The transcription factor ZEB1 regulates stem cell self-renewal and astroglial fate in the adult hippocampus

**DOI:** 10.1101/2020.09.20.305144

**Authors:** B Gupta, AC Errington, S Brabletz, MP Stemmler, T Brabletz, FA Siebzehnrubl

## Abstract

Radial glia-like (RGL) cells persist in the adult mammalian hippocampus where they give rise to new neurons and astrocytes throughout life. Many studies have investigated the process of adult neurogenesis, but factors deciding between neuronal and astroglial fate are incompletely understood. Here, we evaluate the functions of the transcription factor zinc finger E-box binding homeobox 1 (ZEB1) in adult hippocampal RGL cells using a conditional-inducible mouse model. We find that ZEB1 is necessary for self-renewal of active RGL cells as well as for astroglial lineage specification. Genetic deletion of *Zeb1* causes differentiation-coupled depletion of RGL cells resulting in an increase of newborn neurons at the expense of newly generated astrocytes. This is due to a shift towards symmetric cell divisions that consume the RGL cell and generate pro-neuronal progenies. We identify ZEB1 as a regulator of stem cell self-renewal and lineage specification in the adult hippocampus.

## Introduction

The persistence of neural stem cells in the adult hippocampus is well established across many mammalian species (Ming and Song 2011, Gage 2019) including humans (Coras, Siebzehnrubl et al. 2010, Kempermann, Gage et al. 2018). Radial glia-like (RGL) cells within the subgranular zone (SGZ) of the dentate gyrus (DG) reside in a quiescent state, and upon activation are capable of undergoing self-renewal, or differentiating into neurons or astrocytes. Neurogenesis, the process of neuron production, occurs in a step-wise manner during which RGL cells give rise to a group of intermediate progenitor cells (IPCs) (Kempermann, Jessberger et al. 2004). IPCs can clonally expand (Pilz, Bottes et al. 2018) and commit to the neuronal lineage to become neuroblasts, which mature into granule neurons that incorporate into the DG circuitry. While it is well established that RGL cell numbers decrease with age (Encinas, Michurina et al. 2011, Martin-Suarez, Valero et al. 2019), it remains unclear whether this is due to a limited number of cell divisions per RGL cell (Encinas, Michurina et al. 2011), or whether the RGL cell pool is sustained over the lifetime of an animal, with a slight age-related decline (Bonaguidi, Wheeler et al. 2011). Likewise, it is not yet fully established whether astrogliogenesis in the DG occurs through terminal differentiation of RGL cells (Encinas, Michurina et al. 2011), concurrently with neurogenesis (Bonaguidi, Wheeler et al. 2011), through as-yet unidentified astrocyte-specific RGL cells, or through a combination of all three.

Certain transcription factors have been identified that regulate astroglial versus neuronal specification (Bonzano, Crisci et al. 2018, White, Fan et al. 2020), but transcriptional mechanisms underpinning the choice between self-renewal and lineage commitment in the adult brain remain incompletely understood.

Zinc finger E-box binding homeobox 1 (ZEB1) is one of two members of the ZEB family of transcription factors, which are well known regulators of stem cell self-renewal and epithelial-mesenchymal transition (EMT) in solid tissues (Goossens, Vandamme et al. 2017, Stemmler, Eccles et al. 2019). Through these functions, ZEB1 has also been shown to promote malignant growth and dissemination of brain tumours (Edwards, Woolard et al. 2011, Siebzehnrubl, Silver et al. 2013, Rosmaninho, Mukusch et al. 2018). ZEB1 can act as either a transcriptional activator or repressor depending on the recruited cofactors (Spaderna, Schmalhofer et al. 2008, Rosmaninho, Mukusch et al. 2018). ZEB1 expression is crucial for the maintenance of the embryonic radial glial cells in an undifferentiated state, and its downregulation drives the correct maturation and migration of cerebellar and cortical neurons during development (Singh, Howell et al. 2016, Liu, Liu et al. 2019, Wang, Xiao et al. 2019). Zeb1-null mice die perinatally with severe skeletal defects, craniofacial abnormalities, and limb defects, as well as respiratory failure and T cell deficiency; this indicates the importance of ZEB1 in cell polarity (Takagi, Moribe et al. 1998). The lethal phenotype of the Zeb1-null mouse precluded the study of ZEB1 in the adult brain (Takagi, Moribe et al. 1998, Brabletz, Lasierra Losada et al. 2017). However, the recent generation of a conditional *Zeb1* knockout model (Brabletz, Lasierra Losada et al. 2017) has enabled investigating ZEB1 functions beyond early development.

Here, we used the Tamoxifen-inducible GLAST::CreER^T2^ model (Mori, Tanaka et al. 2006, DeCarolis, Mechanic et al. 2013) to investigate the effects of *Zeb1* deletion in adult hippocampal neurogenesis. ZEB1 is expressed in hippocampal RGL and intermediate progenitor cells (IPCs), as well as mature astrocytes, but is downregulated in the neuronal lineage. We found that ZEB1 is necessary for the maintenance of activated RGL cells; loss of *Zeb1* led to a differentiation-coupled depletion of the RGL pool accompanied by a transient increase in neurogenesis and a loss of SGZ-derived astrocytes. In addition to increased neuronal production, we also found increased survival of neurons during their maturation. Analysis of individual RGL cell clones showed that most clones in *Zeb1* deleted animals contained only two neurons and no active RGL cell, whereas clones in control mice contained active RGL cells and a mixture of neurons and astrocytes. Further analysis of mitotic figures revealed that *Zeb1* deletion resulted in a greater number of symmetrical divisions in comparison to wild type mice, leading to their precocious pro-neuronal differentiation. Together, our data demonstrate that ZEB1 is necessary for self-renewal and astroglial lineage specification of hippocampal RGL cells by promoting asymmetrical cell divisions. This further indicates that astroglial fate specification is dependent on asymmetrical divisions in RGL cells.

## Results

ZEB1 is a known regulator of stemness in many solid tissues (Goossens, Vandamme et al. 2017). Recent studies have shown that ZEB1 functions in embryonic neurogenesis (Singh, Howell et al. 2016, Liu, Liu et al. 2019, Wang, Xiao et al. 2019), and we have previously shown that ZEB1 is crucial for the self-renewal of cancer stem cells in glioblastoma (Siebzehnrubl, Silver et al. 2013, Hoang-Minh, Siebzehnrubl et al. 2018, Jimenez-Pascual, Hale et al. 2019). Thus, we hypothesized that ZEB1 would likely execute similar functions in adult neural stem cells. To evaluate the functions of this transcription factor, we first studied the expression of ZEB1 in the adult hippocampus as a paradigm of a well-characterized neurogenic niche.

### Expression of ZEB1 in hippocampal cell types

We assessed the expression of ZEB1 in the adult hippocampus DG (**Fig. 1A**). Co-expression of ZEB1 with cell type-specific markers was evaluated in 12-week old mouse brain tissue sections. Co-staining for ZEB1 and glial fibrillary acidic protein (GFAP, **Fig. 1B**) showed that ZEB1 was abundantly expressed in RGL cells (**Fig. 1B’**), which were identified by expression of (GFAP) as well as a radial morphology with the soma located in the SGZ (Seri, Garcia-Verdugo et al. 2001). We also detected ZEB1 in mature GFAP+ astrocytes in the molecular layer (ML, **Fig. 1B’’**). We observed ZEB1 expression in virtually all GFAP+ cells. ZEB1 was also present in all proliferation-competent cells within the SGZ, as identified by the proliferation marker MCM2 (**Fig. 1C**). By contrast, ZEB1 was absent within the neuronal lineage, including DCX positive early neuronal cells (Brown, Couillard-Despres et al. 2003), and NeuN positive mature neurons (**Fig. 1D-E**). Hence, ZEB1 is present in RGL cells (GFAP+MCM2- and GFAP+MCM2+), IPCs (GFAP-MCM2+), and in astrocytes, but is downregulated once cells undergo neuronal lineage commitment. This is further supported by published datasets from single cell RNA-sequencing studies (**Fig. S1**; (Hochgerner, Zeisel et al. 2018)). This expression pattern supports a functional role for ZEB1 in adult neural stem and progenitor cells. Interestingly, the continued expression of ZEB1 in astrocytes also suggests a more general role for glial identity, comparable to e.g. SOX2 (Bani-Yaghoub, Tremblay et al. 2006).

**Figure 1:**
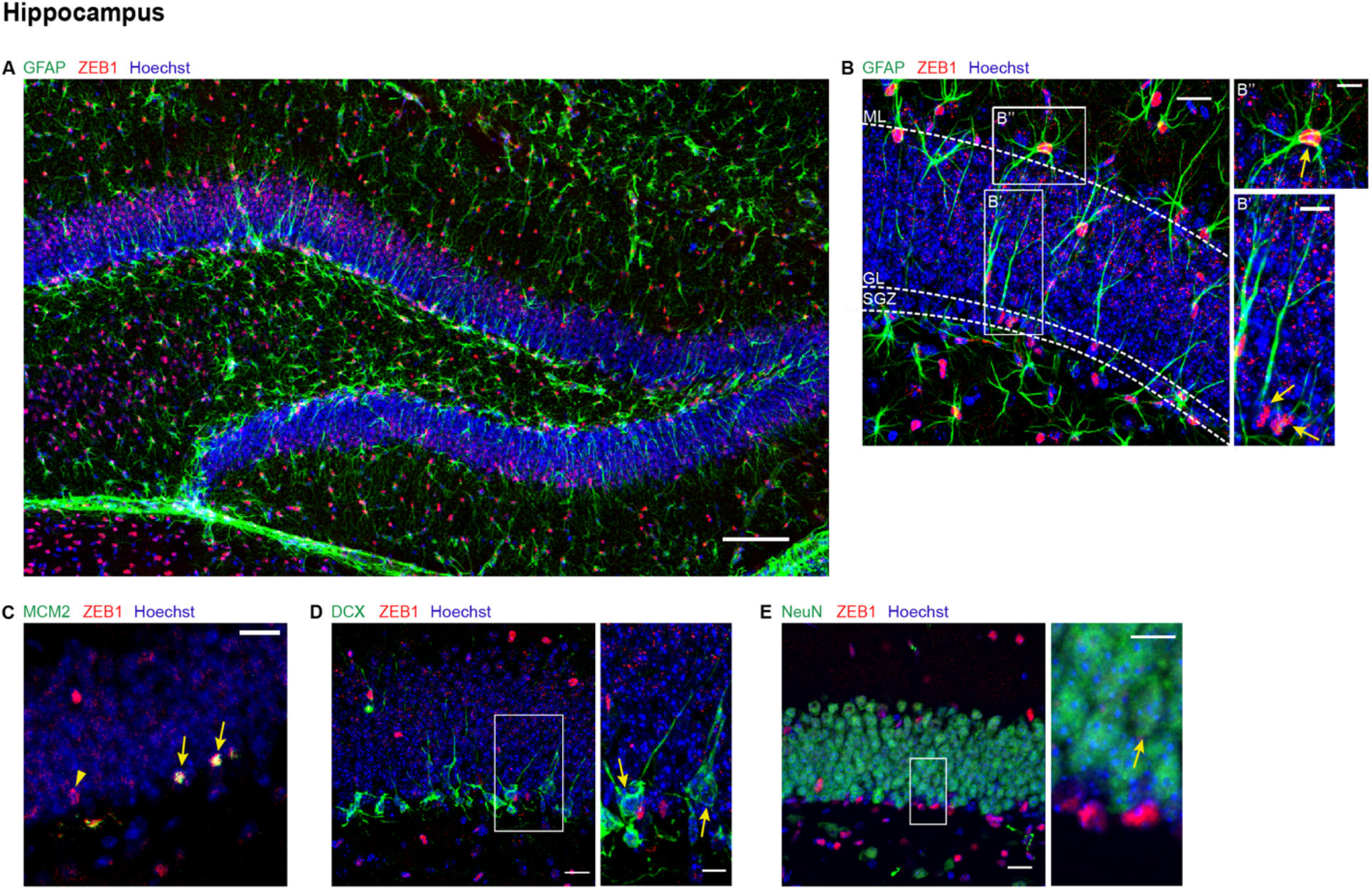
Expression of ZEB1 in the adult mouse hippocampus. **(A)** Overview of whole DG with co-staining of GFAP and ZEB1. **(B)** GFAP and ZEB1 are co-expressed in RGL cells located in the SGZ with a projection crossing the granule layer into the molecular layer (GL; **B’**) and in DG-associated astrocytes in the molecular layer (ML; **B’’**). **(C)** The majority of ZEB1+ cells in the SGZ co-express MCM2 (arrow), but a subset of ZEB1+ cells exists that lack this proliferation-competency marker (arrowhead). **(D-E)** ZEB1 is absent in DCX+ neuroblasts (arrows), as well as in NeuN+ granule neurons (arrow). All scale bars 20 μm (10 μm in insets).

### Characterization of a conditional-inducible *Zeb1* deletion model

To evaluate the function of ZEB1 in RGL cells we employed a loss-of-function approach and generated a conditional-inducible mouse model for deletion of *Zeb1* (**Fig. 2A**). For this, we crossed the Zeb1^f/f^ mouse line (Brabletz, Lasierra Losada et al. 2017) with the tamoxifen-inducible GLAST:CreER^T2^ transgenic line (Mori, Tanaka et al. 2006). The resulting mouse line was further crossed with the Rosa26-tdTomato reporter (Madisen, Zwingman et al. 2010). This model, hereafter referred to as Zeb1^-/-^, enabled the deletion of *Zeb1* in neural stem cells and the astroglial lineage combined with lineage tracing. As controls, we used GLAST:CreER^T2^ / Rosa26-tdTomato mice with wild-type levels of ZEB1 expression (hereafter referred to as control). Tamoxifen (TAM) was administered in 4-5 week old mice and brain tissue harvested at different time points post-induction (**Fig. 2B**).

**Figure 2:**
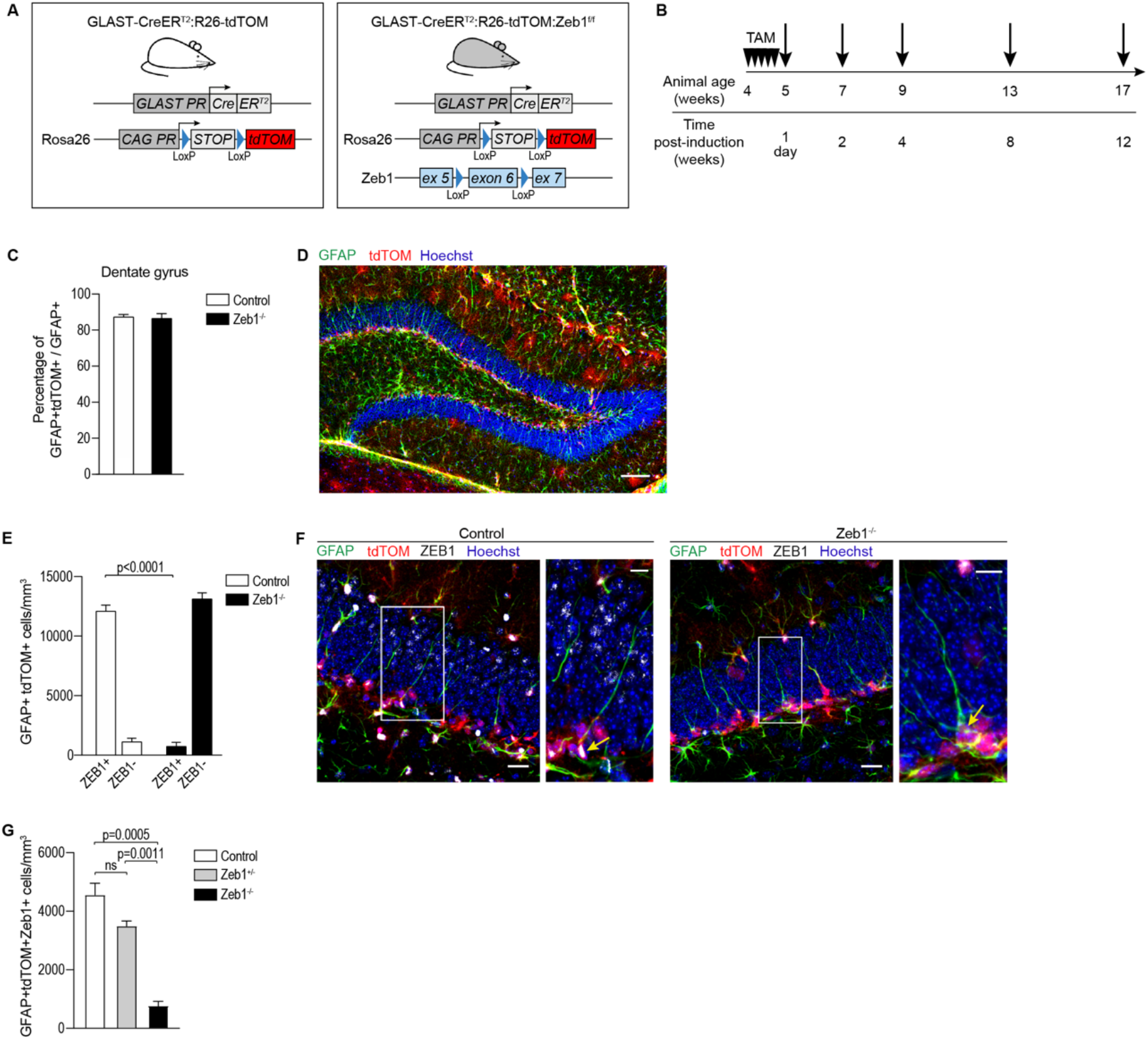
Validation of inducible Zeb1 knockout model. **(A)** Breeding strategy to generate inducible control and Zeb1 knockout mice with endogenous tdTOM reporter expression. **(B)** Mice were administered tamoxifen (TAM) daily for five consecutive days. Tissue was harvested at the indicated time points post-induction (black arrows). **(C)** Induction of tdTOM in GFAP+ cells was comparable in both models. **(D)** Representative image depicting overlap of tdTOM in GFAP+ RGL cells within the SGZ. **(E)** Quantification of ZEB1 expression in GFAP+tdTOM+ RGL cells in the SGZ of control and Zeb1^-/-^ mice 1 day post-induction. **(F)** Representative images showing RGL cells express ZEB1 in control mice, whereas ZEB1 is lost in RGL cells following TAM administration in Zeb1^-/-^ mice. **(G)** Comparison of ZEB1-expressing RGL cells in control, Zeb1^+/-^, and Zeb1^-/-^ mice revealed a significant phenotype only in the full knockout (t-test). All numerical data shown as mean ± SEM. All scale bars 20 μm (10 μm in insets).

We tested the recombination efficiency in control and Zeb1^-/-^ mice by calculating the percentage of GFAP+ RGL cells that co-expressed the endogenous reporter tdTomato (tdTOM) one day after the last TAM injection. Recombination occurred at a high level and to a comparable extent in both models (**Fig. 2C**,**D**).

Next, we quantified the numbers of ZEB1+ and ZEB1-cells in GFAP+tdTOM+ cells one day post-induction to determine the efficiency of Zeb1 deletion in RGL cells. There was a 17-fold decrease in the number of ZEB1+ RGL cells following TAM administration in Zeb1^-/-^ mice compared to controls (**Fig. 2E**,**F**). This validates successful and efficient knockout of *Zeb1* in RGL cells following TAM administration.

Comparison of control and Zeb1^-/-^ mice with compound heterozygotes demonstrated that the phenotype of ZEB1^+/-^ is not significantly different from the controls (**Fig. 2G**). Hence, Zeb1^-/-^ mice can be used to ablate *Zeb1* in DG RGL cells, and bi-allelic deletion of *Zeb1* is needed to obtain a significant phenotype.

### *Zeb1* loss causes depletion of hippocampal RGL cells

Having established that ZEB1 is absent in Zeb1^-/-^ mice immediately after TAM administration, we investigated the longer-term effects of *Zeb1* deletion in hippocampal RGL cells. We quantified quiescent (GFAP+MCM2-; arrowheads) and activated (GFAP+MCM2+; arrows) RGL cells at 1 day (**Fig. 3A**) and 4 weeks (**Fig. 3B**) post-*Zeb1* deletion. The number of quiescent RGL cells was comparable between control and Zeb1^-/-^ mice immediately after induction but was significantly lower by 4 weeks (**Fig. 3C**). By contrast, activated RGL cells showed an immediate decrement that became more pronounced by 4 weeks (**Fig. 3D**). This indicates that *Zeb1* loss causes a steady decline of activated RGL cells, which in turn results in continued recruitment of quiescent RGL cells that ultimately exhausts the hippocampal stem cell pool.

**Figure 3:**
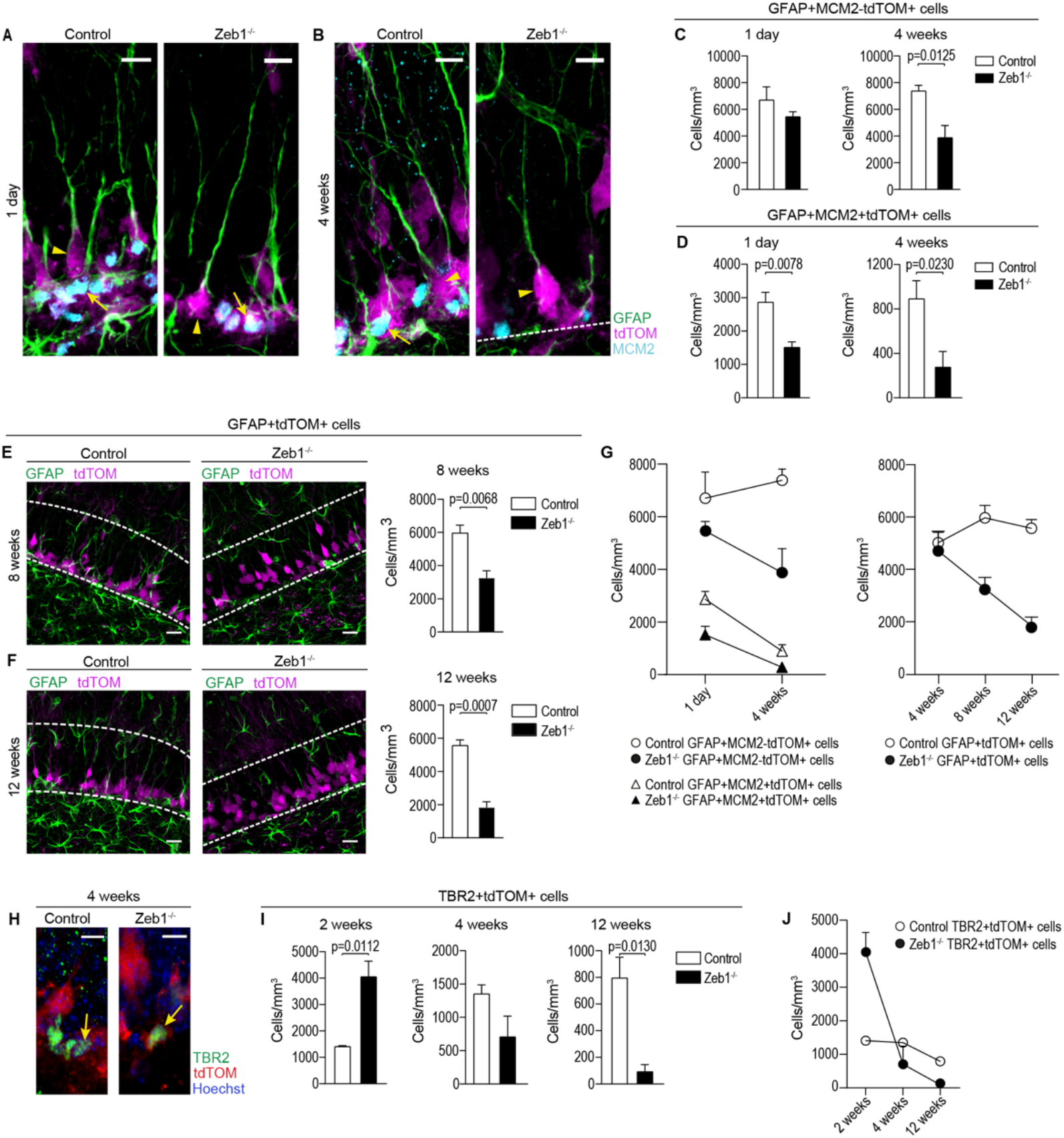
Effects of Zeb1 loss in RGL cells and IPCs. Representative images of quiescent (GFAP+MCM2-tdTOM+; arrowheads) and activated (GFAP+MCM2+tdTOM+; arrow) RGL cells in the SGZ of control and Zeb1^-/-^ mice at 1 day **(A)** and 4 weeks **(B)** post-tamoxifen treatment. **(C)** Numbers of quiescent RGL cells initially remain unchanged, but slowly decline over 4 weeks post-*Zeb1* deletion. **(D)** Contrastingly, numbers of activated RGL cells immediately decrease after *Zeb1* loss in the Zeb1^-/-^ mice and this difference became more pronounced at 4 weeks. **(E)** Representative images and quantification of RGL cells at 8 weeks post-induction showed a continued reduction in the number of RGL cells in Zeb1^-/-^ mice. **(F)** At 12 weeks, representative images and quantification confirmed the progressive long-term depletion of RGL cells in the Zeb1^-/-^ mice. **(G)** Summary graphs depicting quiescent, activated (left), and total (right) RGL populations in control and Zeb1^-/-^ mice. **(H)** Representative images at 4 weeks post-induction and quantification **(I)** of TBR2+ IPCs demonstrate an initial increase in IPCs detected up to 2 weeks post-Zeb1 deletion, followed by a steady decline over the following 10 weeks. **(J)** Summary graph outlining the temporal changes of TBR2+ IPCs. Dashed lines in images demarcate DG boundaries; all scale bars 20 μm (insets: 10 μm).

To confirm this hypothesis, we investigated if the loss of RGL cells continued over time, for which we assessed combined numbers of quiescent and activated RGL cells. Zeb1^-/-^ mice continued to display decreased RGL cell numbers compared to controls by 8 weeks (**Fig. 3E**) and 12 weeks post-induction (**Fig. 3F**). Direct comparison of the trajectories of quiescent, activated, and total RGL cells over time highlights the slow but steady decline of RGL cells in Zeb1^-/-^ mice, while this population remains at steady-state in controls (**Fig. 3G**).

The slow-rate depletion of RGL cells suggests an increased rate of differentiation at the expense of self-renewal in this cell population. We therefore sought to determine whether *Zeb1* loss resulted in altered numbers of hippocampal IPCs by evaluating the number of TBR2+tdTOM+ IPCs (**Fig. 3H**) in control and Zeb1^-/-^ mice between 2 and 12 weeks post-*Zeb1* deletion (**Fig. 3I**). IPC numbers in Zeb1^-/-^ mice were significantly greater at 2 weeks after induction than in controls, but by 4 weeks IPC numbers were comparable in both groups. By 12 weeks post-recombination, this effect was inverted and numbers of TBR2+ cells in Zeb1^-/-^ mice were significantly lower than in controls. These observations are consistent with an increased differentiation of RGL cells causing a transient increase of IPCs that is reverted when RGL cell numbers are depleted (summarized in **Fig. 3J**).

### Loss of *Zeb1* shifts cell fate selection in the hippocampus

We next sought to determine whether the transient IPC increase in Zeb1^-/-^ mice translated into increased numbers of differentiated, lineage-specific progenies. To investigate the fate of the progenitor cells generated through the division-coupled depletion of RGL cells, we assessed the numbers of neuroblasts and neurons, as well as mature astrocytes located within the SGZ between 2- and 12-weeks post-induction.

We first quantified newly generated cells in the neuronal lineage and found that the trajectory of DCX+tdTOM+ neuroblasts in the DG of Zeb1^-/-^ mice followed a similar pattern as IPCs. Compared to controls, the number of DCX+ cells in Zeb1^-/-^ mice was significantly greater at 2 weeks and 4 weeks post-induction. However, this distribution ultimately inverted and at 8 weeks the number of DCX+ cells was significantly lower in Zeb1^-/-^ mice compared to controls, in line with the observed decline in RGL cells and IPCs (**Fig. 4A-C**).

**Figure 4:**
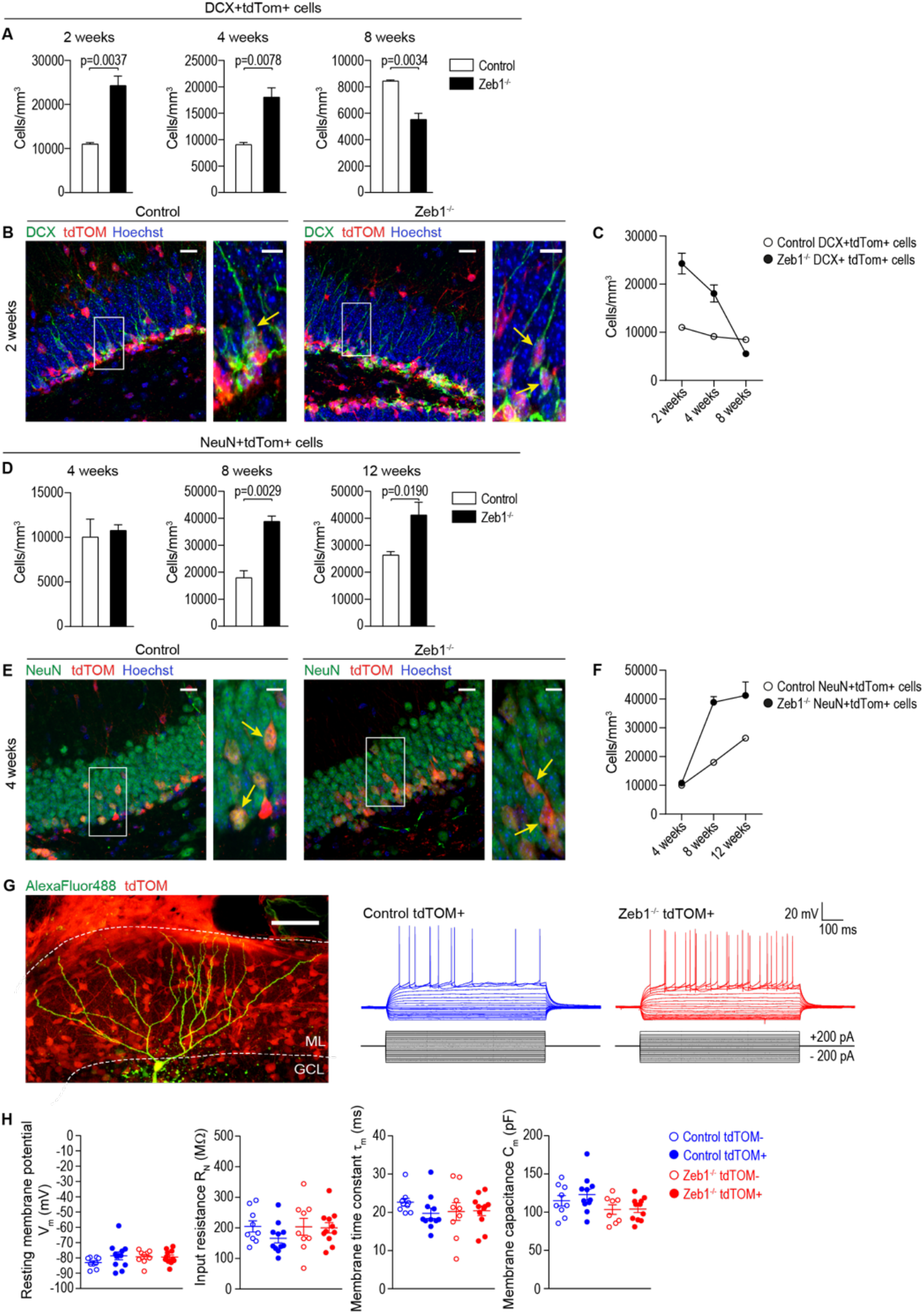
Effects of Zeb1 loss in newborn neurons. **(A)** Quantification of DCX+tdTOM+ neuroblasts in the DG of control and Zeb1^-/-^ mice revealed a two-phased effect: Zeb1 deletion caused an initial increase lasting up to 4 weeks post-induction, with a subsequent decline. **(B)** Representative images of DCX+tdTOM+ neuroblasts at 2 weeks post-induction. **(C)** Summary of neuroblast population dynamics over 8 weeks post-induction in control and Zeb1^-/-^ mice. **(D)** Numbers of NeuN+tdTOM+ granule neurons were comparable between the control and Zeb1^-/-^ mice at 4 weeks post-induction. However, at 8 and 12 weeks post-induction, the number of granule neurons in Zeb1^-/-^ mice was significantly greater than in control mice. **(E)** Representative images at 4 weeks post-induction demonstrate the similar morphology of NeuN+ granule neurons in control and Zeb1^-/-^ mice **(F)** Summary graph showing the increase in granule neurons in the Zeb1^-/-^ mice in comparison to the control mice. **(G)** 2D projection of a 3D 2-photon image stack showing a typical granule cell filled with Alexa488 via the patch clamp recording electrode. Representative recordings from tdTomato expressing Zeb1^-/-^ (red) and control (blue) DGGCs. **(H)** Scatter plots show resting membrane potential (R_m_), input resistance (R_N_), membrane time constant (τ_m_) and membrane capacitance (C_m_) for individual DGGCs overlaid with the mean for each group. Additional graphs and Neurolucida traces are in Fig. S2.

Because of the transient amplification of neuroblasts following *Zeb1* deletion, we probed whether these cells survived the initial stages of maturation and integration to become mature granule neurons. Comparing NeuN+tdTOM+ granule neurons from 4 to 12 weeks post-*Zeb1* deletion, we observed initially no difference in the number of mature neurons between control and Zeb1^-/-^ mice. But at 8- and 12-weeks post-induction the numbers of NeuN+ neurons were significantly greater in Zeb1^-/-^ mice than in controls (**Fig. 4D-F**). Of note, mature neuron numbers showed a linear increase in control mice but appeared to reach a plateau in Zeb1^-/-^ mice between 8 and 12 weeks. Hence *Zeb1* deletion results in a prominent increase in neuronal differentiation and maturation. To test whether newborn neurons displayed normal maturation, we performed patch-clamp electrophysiology (**Fig. 4G**,**H, Fig. S2**). In tdTOM expressing Zeb1^-/-^ and control dentate gyrus granule cells (DGGC) no significant differences were found between resting membrane potential, input resistance, membrane time constant or membrane capacitance 4-5 weeks after TAM injection. We also found no significant differences in these properties between tdTOM-Zeb1^-/-^ and control DGGCs cells or between tdTOM+ and tdTOM-cells of both genotypes (**Fig. 4H**). Furthermore, no differences were observed in cellular excitability with all four groups of DGGCs having similar action potential firing properties (**Fig. S2**). In reconstructed tdTOM+ DGGCs we also found no significant differences in dendritic morphology 4-5 weeks after induction by TAM injection (**Fig. S2**). Interestingly, and in contrast to the effects of *Zeb1* deletion in embryos, our findings suggest that *Zeb1* knockout does not markedly alter the functional and morphological development or migration of DGGC (**Fig. S2**).

In summary, *Zeb1* loss causes depletion of RGL cells by inducing pro-neuronal differentiation, but RGL cells in the DG generate also astrocytes (Bonaguidi, Wheeler et al. 2011, Encinas, Michurina et al. 2011, Gebara, Bonaguidi et al. 2016). We therefore assessed whether the neuronal differentiation-coupled depletion of the RGL cells also affected astrocyte numbers in the SGZ. Because the GLAST promoter is active in both RGL cells and normal astrocytes, we only counted mature astrocytes within the SGZ and excluded astrocytes within other areas of the DG and the hilus. We quantified mature S100β+ astrocytes of control and Zeb1^-/-^ mice from 2 to 12 weeks post-induction. Astrocyte numbers differed not significantly at 2 weeks but showed a progressive decline by 8- and 12-weeks post-induction in Zeb1^-/-^ mice compared to controls (**Fig. 5A-C**). To further elucidate whether *Zeb1* loss results in diminished production of new astrocytes, we analyzed the cellular composition of individual RGL clones.

**Figure 5:**
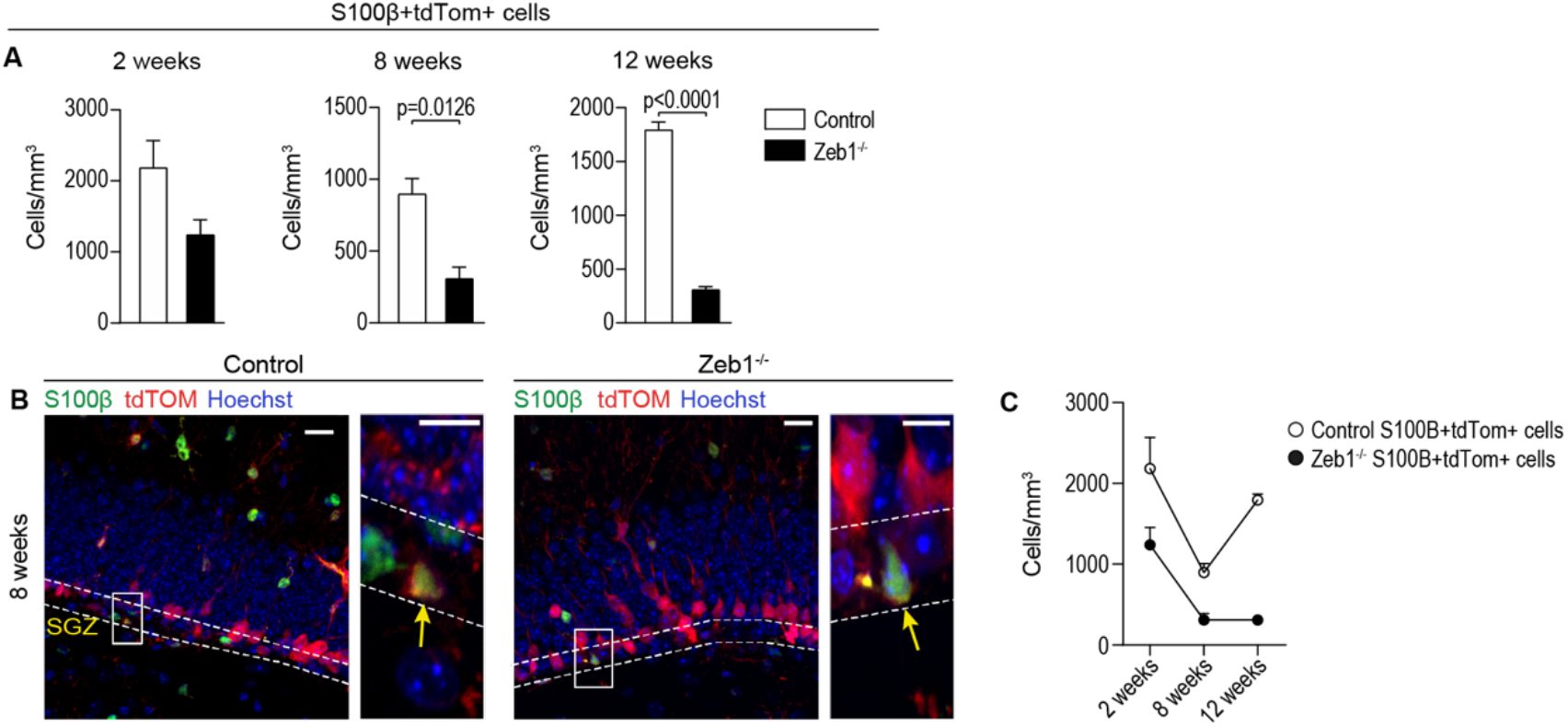
Effects of ZEB1 loss in SGZ-residing astrocytes. **(A)** The number of S100β+tdTOM+ SGZ astrocytes in control and Zeb1^-/-^ mice showed a progressive decline over a 12-week period. **(B)** Representative images at 8 weeks post-induction, identifying SGZ astrocytes (arrows). **(C)** Summary graph showing the gradual decrease in SGZ astrocytes over time. All scale bars 20 μm (insets: 10 μm).

### Analysis clonal lineages in Zeb1^-/-^ and control mice

To investigate how neurogenesis and astrogliogenesis are altered in Zeb1^-/-^ mice, we employed a low-dose induction paradigm that enabled recombination and subsequent lineage tracing in individual RGL cells (**Fig. 6A**,**B** (Bonaguidi, Wheeler et al. 2011)). We analyzed clonal lineages in control and Zeb1^-/-^ mice 4 weeks after recombination and found that a significantly greater number of clones only contained differentiated progenies and lacked an RGL cell in Zeb1^-/-^ mice compared to controls (**Fig. 6C, S3A**). These RGL-depleted clones support our previous observation that RGL cells are lost following *Zeb1* deletion. Conversely, control mice featured more clones containing an RGL cell as well as differentiated progenies (active clones) compared to Zeb1^-/-^ animals. The frequency of clones containing a single RGL cell (quiescent clones) was similar in control and Zeb1^-/-^ mice (**Fig. 6C, S3A**). This indicates that *Zeb1* loss likely does not affect quiescent RGL cells, but its effects only manifest after RGL cells become activated. This notion is further supported by the delayed decline in quiescent RGL cell numbers, whereas activated RGL cell numbers decrease immediately following TAM administration (**Fig. 3**).

**Figure 6:**
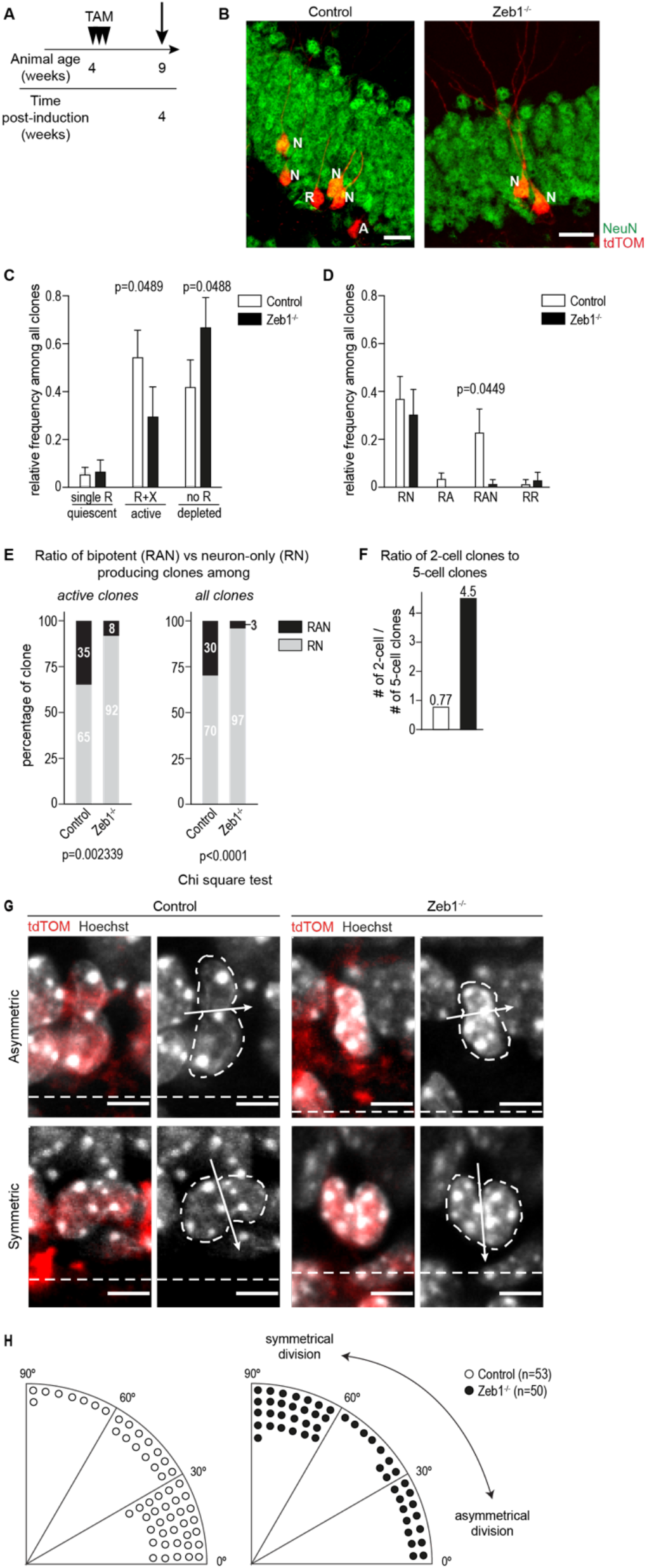
Analysis of RGL cell clones. **(A)** Summary of experimental design. Mice were injected with a low dose of TAM (0.05 mg) and recombination was assessed at 4 weeks post-induction. **(B)** Representative images of RGL cell clones 4 weeks post-induction. **(C)** Relative frequency of quiescent (containing only an RGL cell (single R)), active (containing an RGL cell and any other cell type (R+X)) and depleted (containing only lineage-restricted cells (no R)) clones. Zeb1^-/-^ mice show a significant reduction of active clones and a significant increase in depleted clones compared to controls (n = 38 (control, from 12 hippocampi) vs. 35 (Zeb1^-/-^, from 10 hippocampi). **(D)** Frequencies of active clone subtypes (relative to all clones; n = 26 (control) vs. 14 (Zeb1^-/-^)). Active clones can be either neurogenic (RGL cell and neurons (RN)), astrogliogenic (RGL cell and astrocyte (RA)), bi-lineage (RGL cell, neuron(s) and astrocyte (RAN)), or self-renewing (two RGL cells (RR)). Only RAN clones are significantly reduced in Zeb1^-/-^ mice compared to controls. **(E)** Frequencies of bi-lineage (RAN) versus neuron-only producing (RN) clones across active clones (containing RGL cell) and all clones are significantly lower in Zeb1^-/-^ mice. **(F)** Ratio of clones containing 2 cells versus clones containing 5 cells is 6-fold higher in Zeb1^-/-^ mice compared to controls, indicating a substantially higher number of clones containing 2 cells in this group. **(G)** Representative images of cleavage plane orientation in RGL cells undergoing asymmetric (top) or symmetric (bottom) division in control and Zeb1^-/-^ mice. **(H)** Quantification of RGL cell division angles, binned into 30° groups revealed a significantly higher number of symmetrical divisions in Zeb1^- /-^ mice.

Because of the increase in neurogenesis and concomitant decrease in astrocyte numbers in the SGZ (**Fig. 5**), we suspected that *Zeb1* loss may cause a potential shift in lineage selection. To validate this hypothesis, we analyzed clonal lineages from individual RGL cells in active and depleted clones (Bonaguidi, Wheeler et al. 2011). In Zeb1^-/-^ mice, we found a significantly lower number of clones containing astrocytes compared to controls (**Fig. 6D**). While we found few clones only containing an RGL cell and an astrocyte (RA) in controls, these were absent in Zeb1^-/-^ mice. More importantly, approx. 30% of active clones in control mice contained both astroglia and neurons (RAN), whereas we found only a single bi-lineage clone in Zeb1^-/-^ animals. Of note, there was no significant difference in the frequency of clones that produce only neurons (RN) between controls and Zeb1^-/-^ mice. We next compared the ratio of neuron-only producing versus bi-lineage (neuron and astrocyte producing) clones and found that this ratio is significantly skewed towards neuron-only producing clones after *Zeb1* loss (**Fig. 6E, S3B**,**C**). This indicates that ZEB1 is not only relevant for self-renewal, but also for lineage selection in hippocampal RGL cells.

To elucidate the capacity for clonal expansion following *Zeb1* deletion, we assessed the number of cells per clone in both groups. When comparing the cell numbers across individual clones, we observed an enrichment of clones containing either 2 or 5 cells in control mice, whereas most Zeb1^-/-^ clones contained 1-3 cells (**Fig. S3D**). This is consistent with a premature differentiation of RGL cells in Zeb1^-/-^ mice, which prohibited further clonal expansion. In line with this, we found that most neurons in control mice were included in active clones (**Fig. S3E**), whereas in Zeb1^-/-^ mice numbers of neurons were similar in active and depleted clones (**Fig. S3F**). Premature differentiation of RGL cells after *Zeb1* loss should result in an increased frequency of smaller clones. Therefore, we analyzed the ratio of clones containing 2 cells versus clones containing 5 cells and found that this is close to 1 in controls (i.e. similar numbers of clones contain 2 or 5 cells), but in Zeb1^-/-^ clones this ratio was almost 6-fold higher (i.e. clones containing 2 cells are 6-fold enriched; **Fig. 6F**). This further supports that *Zeb1* deletion causes loss of RGL cells due to premature differentiation.

We noted that this preferential production of 2-cell clones that lack an RGL cell but contain 2 neurons (**Fig. S3G)** is suggestive of symmetrical cell division that causes differentiation of the mother RGL cell. Therefore, we measured cleavage plane orientation of dividing RGL cells relative to their orientation along the SGZ axis in control and Zeb1^-/-^ mice (**Fig. 6G**,**H**). We grouped cleavage plane angles into 30° bins and found that *Zeb1* loss results in a significant shift in cell division angles. While in control animals most RGL cell divisions occurred along the horizontal axis (i.e. asymmetrical (Knoblich 2008)), the division plane was mostly vertical in Zeb1^-/-^ mice. This indicates that Zeb1^-/-^ RGL cells are more likely to undergo symmetrical division and, taken together with the greater probability for Zeb1^-/-^ clones to contain 2 neurons, supports that these symmetrical divisions are neurogenic and cause depletion of the RGL cell.

### No change in proliferation but increased neuronal survival in Zeb1^-/-^ mice

Our data support that the increase in newborn neurons following *Zeb1* deletion is driven by symmetrical divisions of RGL cells that generate neuroblasts at the expense of RGL cell maintenance. Thus, numbers of RGL cells decline, while neuron numbers increase as long as additional RGL cells can be recruited. In the ventricular zone of the developing embryo, Zeb1 deficiency results in diminished proliferation of radial glia (Liu, Liu et al. 2019). Thus, it is conceivable that RGL cells may be dividing slower after Zeb1 deletion, contributing to the smaller clonal size observed in the former group.

To determine whether proliferation rates of stem/progenitor cells differed between Zeb1^-/-^ and control mice, we first quantified Ki67+tdTOM+ progenitor cells from 1 day to 12 weeks post-induction. There was no significant difference in proliferating cells between Zeb1^-/-^ and control at any timepoint (**Fig. 7A**,**B**). Ki67 is expressed during G1, S and G2 phases of the cell cycle and therefore quantification of proliferating cells based on Ki67 alone can be skewed (Miller, Min et al. 2018).

**Figure 7:**
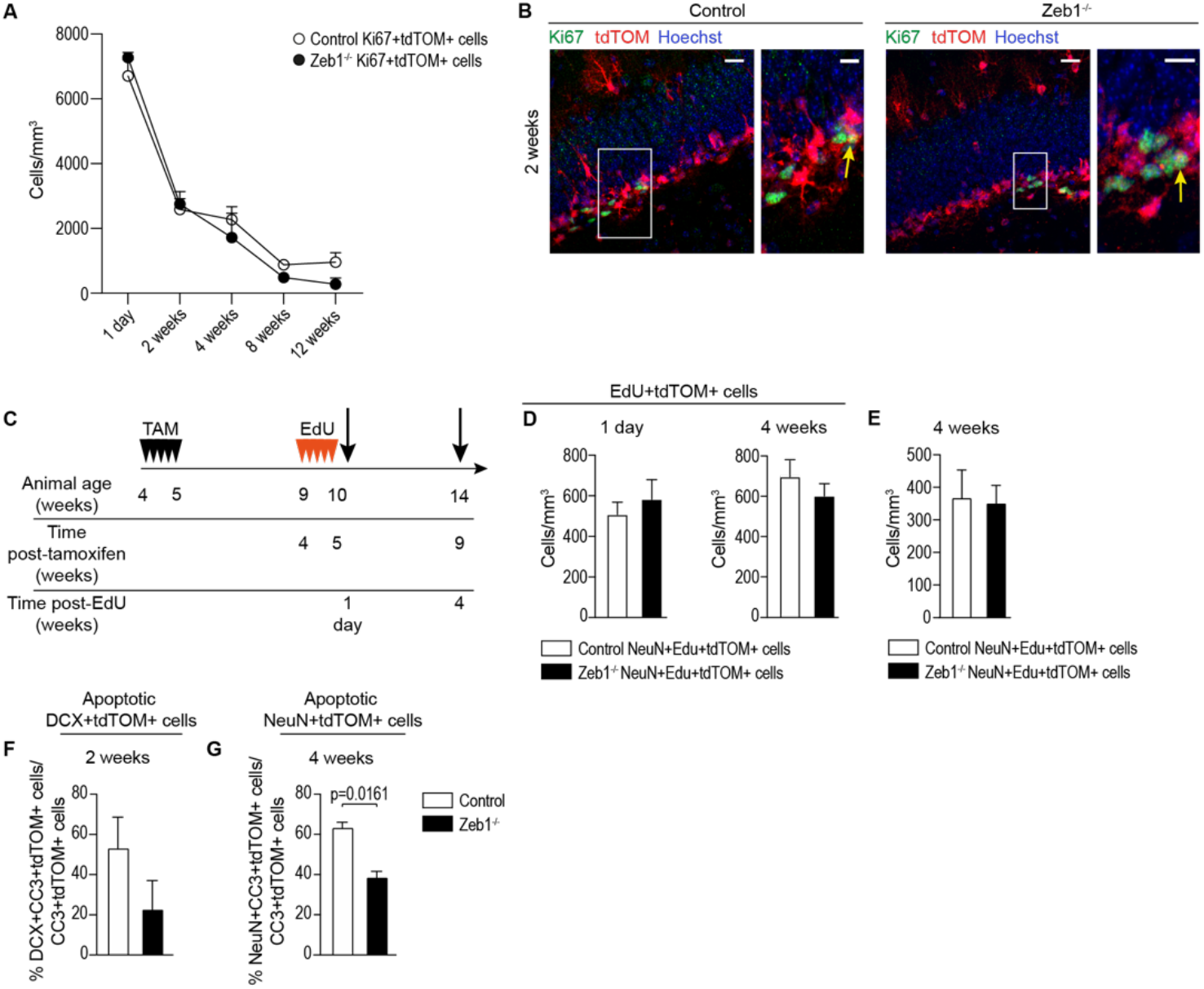
*Zeb1* deletion does not result in slower proliferation rates but increases neuronal survival. **(A)** Quantification of proliferating Ki67+ cells in control and Zeb1^-/-^ mice. Both genotypes show a gradual decrease in proliferating cells over 12 weeks post-induction with no significant differences at any time point. **(B)** Representative images of Ki67+ cells (arrows) used for cell quantification at 2 weeks post-induction. **(C)** Schematic of EdU administration at 4 weeks post-tamoxifen (PT); tissue harvesting was carried out at 1 day and 4 weeks post-EdU injections. **(D)** No difference was observed in the number of EdU+tdTOM+ cells in the control and Zeb1^-/-^ mice at 1 day and 4 weeks post-EdU injections. **(E)** Numbers of EdU-labelled tdTOM+ neurons at 4 weeks PE was comparable between the control and Zeb1^-/-^ cohorts. **(F-G)** Quantification of the fraction of apoptotic cleaved caspase-3+ neuroblasts (DCX+, **F**) and neurons (NeuN+, **G**) demonstrated a decrease in the number of apoptotic neurons in the Zeb1^-/-^ mice in comparison to controls at 4 weeks post-induction.

To further elucidate potential changes in proliferation, we performed EdU labelling *in vivo* 4 weeks post-TAM administration (**Fig. 7C**). We then analyzed numbers of EdU+tdTOM+ cells at 1 day post-EdU but did not find significant differences between control and Zeb1^-/-^ mice (**Fig. 7D**). This supports that cell proliferation rates do not differ between controls and Zeb1^-/-^ mice. To investigate the long-term progenies of dividing cells, we quantified EdU+tdTOM+ (**Fig. 7D**) and the numbers of NeuN+ neurons among these (**Fig. 7E**) at 4 weeks post-EdU. There was no significant difference in long-term EdU-labelled cell numbers or mature neurons derived from EdU-incorporating cells. It is important to note, however, that the numbers of active RGL cells in Zeb1^-/-^ mice at the timepoint of labelling are only about half that of controls (**Fig. 3D**). Therefore, comparable numbers of EdU+NeuN+ neurons were generated by half the number of Zeb1^-/-^ RGL cells, supporting that RGL cells in Zeb1^-/-^ mice undergo symmetric divisions producing two progenies committed to the neuronal lineage.

Most newborn neurons generated during neurogenesis undergo apoptosis and only a small fraction survives and integrates successfully into the hippocampal circuitry (Dayer, Ford et al. 2003, Ryu, Hong et al. 2016). Changes in neuronal survival may considerably impact the number of neurons generated by neurogenesis (Sierra, Encinas et al. 2010). Therefore, we determined whether *Zeb1* loss affected survival of neuronal progenies and quantified co-expression of the apoptotic marker cleaved caspase 3 with either DCX (at 2 weeks post-induction) or NeuN (at 4 weeks post-induction). The number of apoptotic DCX+ neuroblasts was not significantly different between *Zeb1* deleted mice and controls but we did find significantly fewer apoptotic NeuN+ granule neurons in the Zeb1^-/-^ DG (**Fig. 7F**,**G**). This indicates that *Zeb1* loss promotes long-term survival of newly generated hippocampal neurons, which may contribute to the increased levels of neurogenesis following *Zeb1* deletion.

In summary, our data support that *Zeb1* deletion in hippocampal RGL cells promotes their differentiation into neuronal progenies, while simultaneously preventing astroglial fate. Zeb1^-/-^ RGL cells preferentially undergo symmetrical divisions with both daughter cells committed to the neuronal lineage. This effectively eliminates the mother RGL cell and prevents generation of astrocytes. Newborn Zeb1^-/-^ neurons have a higher survival rate than control neurons and integrate into the DG.

## Discussion

The transcription factor ZEB1 was historically most prominently associated with epithelial-mesenchymal transition (Peinado, Olmeda et al. 2007). While primary EMT is an essential mediator of cell state transitions in development and essential for e.g. neural crest formation (Acloque, Adams et al. 2009), the relevance of this process during homeostasis of non-epithelial tissues, such as the adult brain, remains unclear. More recently, the role of EMT-associated transcription factors in the maintenance of stem cell phenotypes have come into focus, prompting a call for re-evaluating their functions ‘beyond EMT and MET’ (Goossens, Vandamme et al. 2017). In line with this notion, ZEB1 is a key mediator of cancer stemness in glioblastoma (Siebzehnrubl, Silver et al. 2013, Rosmaninho, Mukusch et al. 2018), where it is part of an autoregulatory transcription factor loop together with SOX2 and OLIG2 (Singh, Kollipara et al. 2017), two transcription factors with well-established functions in neural stem/progenitor cells (Ligon, Huillard et al. 2007, Suh, Consiglio et al. 2007). Here, we investigated the consequences of *Zeb1* deletion in adult neural stem/progenitor cells. In our model, conditional-inducible deletion of *Zeb1* resulted in rapid and sustained loss of ZEB1 within the DG that was apparent as early as 1 day following TAM administration.

Constitutive deletion of *Zeb1* is lethal around birth and conditional-inducible models have been developed only recently (Brabletz, Lasierra Losada et al. 2017). Therefore, the effects of *Zeb1* deletion in the brain have so far only been investigated during embryonic development. Two studies have reported that downregulation of ZEB1 expression is necessary for the neuronal commitment of precursor cells, allowing them to gain a neuronal identity while migrating to their maturation destinations in the developing cerebellum and cortex, respectively (Singh, Howell et al. 2016, Wang, Xiao et al. 2019). This is partially mirrored in the adult hippocampus as we observed a lack of ZEB1 in the neuroblast and granule neuron populations. Consequently, *Zeb1* deletion results in increased neurogenesis in both embryo and postnatal development.

Studies investigating ZEB1 functions in embryonic neurodevelopment have shown that *Zeb1* loss results in aberrant neuronal morphology and positioning (Singh, Howell et al. 2016, Liu, Liu et al. 2019). By contrast, we found that morphology and migration of adult-born granule neurons in the hippocampus are not significantly altered following *Zeb1* deletion. It is conceivable that adult hippocampal neurogenesis differs from cortical neurogenesis and that hippocampal granule neurons do not need ZEB1 to mature. Alternatively, migration distances in the adult hippocampus may be too short for a noticeable effect on cell migration.

RGL cells constitute resident stem cells of the hippocampal DG and are capable of generating both neurons and astrocytes throughout life (Palmer, Takahashi et al. 1997, Seri, Garcia-Verdugo et al. 2001, Bonaguidi, Wheeler et al. 2011, Encinas, Michurina et al. 2011). We report a steady loss of RGL cells following *Zeb1* deletion, and our data indicate that this depletion is coupled with differentiation, and not cell death. This confirms that ZEB1 is necessary for stem cell maintenance through self-renewal in the adult CNS. The overall increase in granule neuron output over the investigated 12-week period demonstrates that instead of immediately undergoing apoptosis following the deletion of *Zeb1*, the RGL cells gave rise to neuronal progenies. The slow depletion of RGL cells over time indicates a differentiation-coupled exhaustion of the RGL cell pool (Gao, Ure et al. 2011). This RGL cell differentiation is linked to cell division, because most RGL cell-depleted clones contained more than one neuron, indicating that the labelled RGL cells divided at least once prior to depletion. We found a gradual decrease that initially only affected activated RGL cells (GFAP+ MCM2+), whereas numbers of quiescent RGL cells (GFAP+ MCM2-) declined with a 4-week delay. Therefore, *Zeb1* loss does not immediately affect quiescent RGL cells (despite their expression of ZEB1) and numbers of quiescent RGL cells likely only decrease when they are recruited to replenish lost activated RGL cells. This is further supported by our analysis of individual RGL cell clones, which showed comparable numbers of quiescent RGL clones between control and Zeb1^-/-^ mice, although the overall number of quiescent RGL clones was low (3 vs. 4). The existence of a third group of ‘resilient’ RGL cells has been suggested (Ziebell, Martin-Villalba et al. 2014), which may act as an additional reservoir that can replenish activated RGL cells. Resilient RGL cells may contribute to the slow rate of exhaustion of the quiescent RGL cells observed in our study, but because all GFAP+ cells with radial morphology express ZEB1, we cannot distinguish between resilient and quiescent RGL cells here.

During adult neurogenesis, quiescent RGL cells become activated to generate new neurons. While some questions remain, there is consensus that RGL cells undergo a limited number of divisions (most likely three) after which they either revert back to quiescence or terminally differentiate (Bonaguidi, Wheeler et al. 2011, Encinas, Michurina et al. 2011, Lazutkin, Podgorny et al. 2019). In this context it is interesting to note that the ratios of RGL cell-containing to RGL cell-depleted clones is approximately 3:1 in control mice, suggesting that over the 4-week chase period 1 out of 4 clones differentiated. Following *Zeb1* deletion, we find an even ratio of RGL-depleted to RGL-containing clones (1:1), indicating that Zeb1^-/-^ RGL cells are more likely to terminally differentiate. Furthermore, most 2 cell clones in the *Zeb1*-deficient DG were depleted and contained 2 neurons, in comparison to controls where the majority of 2-cell clones contained an RGL cell and one non-RGL cell. While the total number of cells and neurons per clone varied widely, most clones in the control mice contained 5 cells, similar to a recent study tracking neurogenesis by live-cell imaging (Pilz, Bottes et al. 2018). A 2-cell clone is consistent with a single division, whereas a 5-cell clone requires multiple divisions. Thus, the higher ratio of 2-cell clones to 5-cell clones in the Zeb1^-/-^ mice is a strong indicator that Zeb1^-/-^ RGL cells only divide once. Because most 2-cell clones in *Zeb1* deleted mice contain 2 neurons and no RGL cell, the most likely scenario is that the RGL cell differentiates into 2 neuronal progenies. While it is conceivable that the higher frequency of 2-cell clones in *Zeb1* deleted mice could be due to lower proliferation of RGL cells or increased apoptosis of their progenies, our quantification of Ki67 and EdU incorporation as well as cleaved caspase-3 argues against this.

Previous studies that have investigated genetic deletion of stem cell transcription factors have reported similar findings on neurogenesis. Deletion RBPJκ, the main effector of Notch signaling, in SOX2-expressing precursors resulted in precocious neuronal differentiation, with an initial increase in the neuronal population alongside the depletion of the precursor pool (Ehm, Goritz et al. 2010). Similarly, REST deficiency in RGL cells leads to a transient increase in neurogenesis, with a simultaneous decrease in RGL cells (Gao, Ure et al. 2011). Loss of *Pax6* results in a smaller GFAP+ RGL cell pool in comparison to wild type, with thinner and undeveloped radial processes, coupled with abnormal neuronal progenitors, indicating impaired neurogenesis (Maekawa, Takashima et al. 2005). Deletion of COUP-TFI results in an increase in astrogliogenesis at the expense of neurogenesis, but no loss of RGL cells was observed (Bonzano, Crisci et al. 2018). Because of the link between ZEB1 and SOX2 in glioblastoma (Singh, Kollipara et al. 2017), it is noteworthy to compare the functions of both transcription factors in adult neurogenesis. Interestingly, loss of *Sox2* decreased the RGL cell pool by over 40% alongside decreased cell proliferation but had only limited effects on neurogenesis (Favaro, Valotta et al. 2009). This may be due to a cell context-dependent function of SOX2, with SOX2 inhibiting the expression of NeuroD1 in RGL cells, while inducing NeuroD1 when expressed in IPCs (Kuwabara, Hsieh et al. 2009). Here, we show that *Zeb1* loss causes differentiation of RGL cells into the neuronal lineage, thus inducing a transient increase in neurogenesis. The increase in neuroblast production is further amplified by decreased neuronal apoptosis during maturation, thus elevated neuron numbers are a result of both production and survival. Mature Zeb1^-/-^ neurons show electrophysiological and morphological characteristics of normal hippocampal granule cells.

RGL cells have been found to produce both neurons and astrocytes (Steiner, Kronenberg et al. 2004, Bonaguidi, Wheeler et al. 2011, Gebara, Bonaguidi et al. 2016, Bonzano, Crisci et al. 2018), but whether a dedicated astrocyte-specific stem cell exists as well is not yet resolved. A previous study concluded that RGL cells terminally differentiate into astrocytes after three division cycles (Encinas, Michurina et al. 2011), while another report found that RGL cells can produce astrocytes during any of their divisions (Bonaguidi, Wheeler et al. 2011). Both of these studies used the Nestin::Cre line to drive reporter expression. A third study, using the Ascl1::Cre driver, did not observe astrocyte production from RGL cells (Pilz, Bottes et al. 2018). Here, we used the GLAST::CreER^T2^ line (Mori, Tanaka et al. 2006) to induce expression of the fluorescent reporter tdTOM in RGL cells and astrocytes. We cannot fully exclude that quantified SGZ astrocytes may have been derived from other astrocytes and not RGL cells, but the proximity of quantified astrocytes to neighboring cells within a clone combined with a lack of other astrocytes in the vicinity argues against this. We found a combination of clones producing only neurons, only astrocytes, and both lineages, similar to previous reports (Bonaguidi, Wheeler et al. 2011, Gebara, Bonaguidi et al. 2016). In the present study, the ratio of clonal lineages among active clones in control mice was approximately 1:4:8 (astrocyte-specific : bi-lineage : neuron-specific), while Bonaguidi et al. observed a ratio of approx. 2:1:3 (astrocyte-specific : bi-lineage : neuron-specific). It is noteworthy that we induced recombination in 4-5-week-old mice, while most previous studies induced recombination in 8-9-week-old mice. Astrocyte production in the hippocampus increases with age (Encinas, Michurina et al. 2011) which may account for the slightly lower ratios of astrocyte-producing clones in our study. Additionally, different Cre-driver lines used (e.g. Nestin::Cre vs. GLAST::Cre) may also affect clonal lineage distribution. The fact that others and we observed the existence of bi-lineage clones argues against an astrocyte-restricted precursor cell, although it is conceivable that RGL cells are heterogeneous with some cells restricted to a particular lineage (astrocytic or neuronal), and others being more potent. While we observed clones that produced only one astrocyte and no neuronal progenies, we cannot exclude that these may have produced neurons that died before detection. Because we followed clonal production over 4 weeks, it is further possible that uni-lineage clones may produce cells from additional lineages over longer time periods.

In the embryonic CNS, a series of symmetrical divisions that expand the RG cell pool precede a switch to neurogenic asymmetrical divisions, followed by gliogenic asymmetrical divisions (Falk and Gotz 2017, Allen and Lyons 2018). Here, we report that *Zeb1* loss in the adult hippocampus promotes symmetrical RGL cell divisions tied to the production of neuronal lineage-committed progenitors, resulting in a premature depletion of RGL cells. Thus, ZEB1 functions to preserve the RGL cell pool in the adult DG through asymmetrical divisions. Notably, ZEB1 is asymmetrically distributed during cell division in precancerous adenomas (Liu, Siles et al. 2018). It is therefore tempting to speculate that ZEB1 may control its own distribution during mitosis.

## Materials and Methods

### Animals and tamoxifen administration

Animal experimentation was approved by and carried out in accordance with regulations of the UK Home Office. Both male and female mice were used for all experiments. All animals were housed in a 12 h light/dark cycle and given access to food and water *ad libitum*. A transgenic mouse line with loxP sites flanking exon 6 of the Zeb1 gene (Brabletz, Lasierra Losada et al. 2017), was crossed with the GLAST::CreER^T2^ mouse line (kind gift from M. Götz, Munich; (Mori, Tanaka et al. 2006)) and further crossbred with the Rosa26^lox-stop-lox-tdTomato^ strain (kind gift from O. Sansom, Glasgow; (Madisen, Zwingman et al. 2010)). GLAST::CreER^T2^-Rosa26^lox-stop-lox-tdTomato^ mice with wild-type ZEB1 expression were used as controls (referred to as the control strain). For regular transgene induction, a 2 mg dose of tamoxifen (Sigma) prepared in corn oil was injected i.p. into 4-5-week-old mice daily for five consecutive days (Jahn, Kasakow et al. 2018). For clonal analysis, a 50 μg dose of tamoxifen in corn oil was injected i.p. into 4-5 week-old mice daily for three consecutive days. Mice were transcardially perfused and the brains were harvested for histological analysis at timepoints indicated in the text.

### Tissue processing, immunostaining, and confocal imaging

Tissue was processed as previously described (Jimenez-Pascual, Hale et al. 2019). Briefly, 30 μm thick coronal sections of the dorsal hippocampus were maintained in serial order and analyzed at a median bregma -1.8 mm following immunofluorescence staining using confocal imaging on a Zeiss LSM710 microscope with ZEN software. Primary and secondary antibodies used in this study are provided in **Table S2**. An average of four images was obtained spanning the length of the suprapyramidal blade and cells were counted using ImageJ v1.52K. Data represent n=3 mice per genotype, with a minimum of two sections analyzed per animal. The primary researcher responsible for quantification was not blinded during the counting process, but a secondary blinded researcher confirmed an average of one set of technical replicate counts per cell marker per group.

### 5-ethynyl-2’deoxyuridine (EdU) treatment

The EdU Click assay (Sigma) was used to detect proliferating cells and carry out lineage tracing *in vivo*. Four weeks following TAM administration, mice were i.p. injected with 50 mg/kg EdU for five consecutive days. Mice were transcardially perfused and the brains were harvested for histological analysis at timepoints indicated in the text and processed for EdU detection according to the manufacturer’s protocol.

### Clonal analysis

Analysis of individual clones was carried out as described in (Bonaguidi, Wheeler et al. 2011). Cells belonging to a clone were identified by proximity, residing within a 90 μm radius. A combination of fluorescent markers and cell morphology was used to identify DG cell types and is presented in **Table S3**.

### Cleavage plane measurements

For quantification of cell division angles, mitotic figures of tdTOM+ cells with RGL morphology were assessed in sections containing the SGZ stained with Hoechst from 7-8 different mice per genotype. ImageJ was used to quantify the cell cleavage angle by drawing a line along the cleavage furrow. A line drawn along the interface between the hilus and SGZ of the DG was used as reference for cleavage plane angles. Subsequently, the angle measurements were binned into three categories: horizontal (0-30°), intermediate (30-60°), and vertical (60-90°).

### Brain slice preparation, electrophysiology and 2-photon imaging

Brain slice preparation and electrophysiology was performed as described previously (Trent, Hall et al. 2019). Animals of either sex were deeply anaesthetised using isoflurane, decapitated and their brains removed into chilled (1-3°C) cutting solution containing (in mM) 60 sucrose, 85 Nacl, 2.5 KCl, 1 CaCl_2_, 2 MgCl_2_, 1.25 NaH_2_PO_4_, 25 NaHCO_3_, 25 D-glucose, 3 kynurenic acid, 0.045 indomethacin. Horizontal hippocampal brain slices (300 μm) containing the dentate gyrus, prepared from 8 weeks old Zeb1^-/-^ and control mice 4-5 weeks after TAM injection, were initially stored for 20 minutes at 35°C in sucrose-containing solution and subsequently maintained at room temperature in artificial CSF (aCSF) containing (in mM) 125 NaCl, 2.5 KCl, 1 CaCl_2_, 1 MgCl_2_, 1.25 NaH_2_PO_4_, 25 NaHCO_3_, 25 D-glucose (305 mOSm) and used within 4-6 hours. For recording, slices were transferred to a submersion chamber continuously perfused with warmed (33-34°C) aCSF containing (in mM) 125 NaCl, 2.5 KCl, 2 CaCl_2_, 1 MgCl_2_, 1.25 NaH_2_PO_4_, 25 NaHCO_3_, 25 D-glucose (305-10 mOSm, pH 7.4) at a flow rate of 3 ml.min^-1^. Electrophysiological recordings were performed on dentate gyrus granule cells (DGGC) and of the dentate gyrus granule cell layer. DGGC were identified using Dodt-contrast video microscopy and TAM induced cells selected by their expression of the red fluorescent protein tdTomato following 2-photon excitation at λ = 900 nm (Prairie Ultima 2-photon microscope, Bruker). Whole-cell current clamp recordings were made using a Multiclamp 700B (Molecular Devices) patch clamp amplifier with patch pipettes with resistances 4–6 MΩ when filled with internal solution containing (in mM) 130 K-gluconate, 20 KCl, 10 HEPES, 0.16 EGTA, 2 Mg-ATP, 2 Na2-ATP, 0.3 Na2-GTP, pH 7.3 (295 mOsm). Somatic series resistance at the start of experiments was between 9-15 MΩ and cells showing changes of R_S_ greater than 20% over the course of the experiment were rejected. Data were sampled at 20-40 kHz and low-pass filtered at 6 kHz. Resting membrane potential (V_m_) was measured as the mean membrane potential during a 100 ms period prior to a hyperpolarizing current injection step averaged across 10-20 sweeps. Input resistance (R_N_) was calculated, according to Ohm’s law, by dividing the magnitude of the voltage change (sampled over 100 ms) at the end of 1 second hyperpolarizing current injection response by the amount of injected current (20 pA). Membrane time constant (τ_m_) was measured by fitting a mono-exponential function to the repolarizing phase of the same 20 pA hyperpolarizing current step. Membrane capacitance (C_m_) was calculated using τ = RC by dividing R_N_ by τ_m_.

Neuronal excitability was measured by comparing current injection evoked action potentials in DGGC in Zeb1^-/-^ and control mice. Action potential amplitude, half-width, voltage threshold, δV/δt and rheobase was measured. In order to compare TAM induced DGGCs to the larger population of DGGCs patch clamp recordings were performed from both tdTomato positive and negative cells.

To compare dendritic morphology between TAM induced tdTomato expressing DGGC in Zeb1^-/-^ and control mice, recorded cells were filled via the recording electrode with Alexa Fluor 488 (100 μM). Stacks of 120-250 2-photon images (512 × 512 pixels) were collected at Z intervals of 1 μm. Soma and dendrites of imaged DGGC were reconstructed post hoc from 3D image projections using the semi-automated tracing tool in Neurolucida 360 (MBF Bioscience). Analysis of dendritic morphology was performed on 3D neuronal reconstructions using Neurolucida Explorer.

## Statistical analysis

Statistical testing was carried out using GraphPad Prism 8. For comparison of two groups, t-tests were used. A one-tailed t-test was performed where we hypothesized a specific effect of *Zeb1* deletion based on previous research. For categorical analyses, a Chi square test was used. Unless otherwise specified, data are presented as mean ± SEM.

## Acknowledgements

The authors would like to thank D. Siebzehnruebl for technical assistance and D. Petrik for critical comments on the manuscript. FAS is funded by MRC grant MR/S 007709/1 and BG is funded by a CRUK PhD studentship.

## Author contributions

BG designed and performed experiments, analyzed data and wrote the manuscript. ACE designed, performed and analyzed electrophysiology and 2-photon imaging experiments and contributed to the manuscript. FAS designed the study, analyzed data and wrote the manuscript. SB, MPS and TB provided essential animal models and contributed to the manuscript.

## Declaration of interests

The authors declare no competing interests.

## Notes

### Competing Interest Statement

The authors have declared no competing interest.

## References

Acloque, H., M. S. Adams, K. Fishwick, M. Bronner-Fraser and M. A. Nieto (2009). “Epithelial-mesenchymal transitions: the importance of changing cell state in development and disease.” J Clin Invest 119(6): 1438–1449.

Allen, N. J. and D. A. Lyons (2018). “Glia as architects of central nervous system formation and function.” Science 362(6411): 181–185.

Bani-Yaghoub, M., R. G. Tremblay, J. X. Lei, D. Zhang, B. Zurakowski, J. K. Sandhu, B. Smith, M. Ribecco-Lutkiewicz, J. Kennedy, P. R. Walker and M. Sikorska (2006). “Role of Sox2 in the development of the mouse neocortex.” Dev Biol 295(1): 52–66.

Bonaguidi, M. A., M. A. Wheeler, J. S. Shapiro, R. P. Stadel, G. J. Sun, G. L. Ming and H. Song (2011). “In vivo clonal analysis reveals self-renewing and multipotent adult neural stem cell characteristics.” Cell 145(7): 1142–1155.

Bonzano, S., I. Crisci, A. Podlesny-Drabiniok, C. Rolando, W. Krezel, M. Studer and S. De Marchis (2018). “Neuron-Astroglia Cell Fate Decision in the Adult Mouse Hippocampal Neurogenic Niche Is Cell-Intrinsically Controlled by COUP-TFI In Vivo.” Cell Rep 24(2): 329–341.

Brabletz, S., M. Lasierra Losada, O. Schmalhofer, J. Mitschke, A. Krebs, T. Brabletz and M.P. Stemmler (2017). “Generation and characterization of mice for conditional inactivation of Zeb1.” Genesis.

Brown, J. P., S. Couillard-Despres, C. M. Cooper-Kuhn, J. Winkler, L. Aigner and H. G. Kuhn (2003). “Transient expression of doublecortin during adult neurogenesis.” J Comp Neurol 467(1): 1–10.

Coras, R., F. A. Siebzehnrubl, E. Pauli, H. B. Huttner, M. Njunting, K. Kobow, C. Villmann, E. Hahnen, W. Neuhuber, D. Weigel, M. Buchfelder, H. Stefan, H. Beck, D. A. Steindler and I. Blumcke (2010). “Low proliferation and differentiation capacities of adult hippocampal stem cells correlate with memory dysfunction in humans.” Brain 133(11): 3359–3372.

Dayer, A. G., A. A. Ford, K. M. Cleaver, M. Yassaee and H. A. Cameron (2003). “Short-term and long-term survival of new neurons in the rat dentate gyrus.” J Comp Neurol 460(4): 563–572.

DeCarolis, N. A., M. Mechanic, D. Petrik, A. Carlton, J. L. Ables, S. Malhotra, R. Bachoo, M. Gotz, D. C. Lagace and A. J. Eisch (2013). “In vivo contribution of nestin- and GLAST-lineage cells to adult hippocampal neurogenesis.” Hippocampus 23(8): 708–719.

Edwards, L. A., K. Woolard, M. J. Son, A. Li, J. Lee, C. Ene, S. A. Mantey, D. Maric, H. Song, G. Belova, R. T. Jensen, W. Zhang and H. A. Fine (2011). “Effect of brain- and tumor-derived connective tissue growth factor on glioma invasion.” Journal of the National Cancer Institute 103(15): 1162–1178.

Ehm, O., C. Goritz, M. Covic, I. Schaffner, T. J. Schwarz, E. Karaca, B. Kempkes, E. Kremmer, F. W. Pfrieger, L. Espinosa, A. Bigas, C. Giachino, V. Taylor, J. Frisen and D. C. Lie (2010). “RBPJkappa-dependent signaling is essential for long-term maintenance of neural stem cells in the adult hippocampus.” J Neurosci 30(41): 13794–13807.

Encinas, J. M., T. V. Michurina, N. Peunova, J. H. Park, J. Tordo, D. A. Peterson, G. Fishell, A. Koulakov and G. Enikolopov (2011). “Division-coupled astrocytic differentiation and age-related depletion of neural stem cells in the adult hippocampus.” Cell Stem Cell 8(5): 566–579.

Falk, S. and M. Gotz (2017). “Glial control of neurogenesis.” Curr Opin Neurobiol 47: 188–195.

Favaro, R., M. Valotta, A. L. Ferri, E. Latorre, J. Mariani, C. Giachino, C. Lancini, V. Tosetti, S. Ottolenghi, V. Taylor and S. K. Nicolis (2009). “Hippocampal development and neural stem cell maintenance require Sox2-dependent regulation of Shh.” Nat Neurosci 12(10): 1248–1256.

Gage, F. H. (2019). “Adult neurogenesis in mammals.” Science 364(6443): 827–828.

Gao, Z., K. Ure, P. Ding, M. Nashaat, L. Yuan, J. Ma, R. E. Hammer and J. Hsieh (2011). “The master negative regulator REST/NRSF controls adult neurogenesis by restraining the neurogenic program in quiescent stem cells.” J Neurosci 31(26): 9772–9786.

Gebara, E., M. A. Bonaguidi, R. Beckervordersandforth, S. Sultan, F. Udry, P. J. Gijs, D. C. Lie, G. L. Ming, H. Song and N. Toni (2016). “Heterogeneity of Radial Glia-Like Cells in the Adult Hippocampus.” Stem Cells 34(4): 997–1010.

Goossens, S., N. Vandamme, P. Van Vlierberghe and G. Berx (2017). “EMT transcription factors in cancer development re-evaluated: Beyond EMT and MET.” Biochim Biophys Acta Rev Cancer 1868(2): 584–591.

Hoang-Minh, L. B., F. A. Siebzehnrubl, C. Yang, S. Suzuki-Hatano, K. Dajac, T. Loche, N. Andrews, M. Schmoll Massari, J. Patel, K. Amin, A. Vuong, A. Jimenez-Pascual, P. Kubilis, T. J. Garrett, C. Moneypenny, C. A. Pacak, J. Huang, E. J. Sayour, D. A. Mitchell, M. R. Sarkisian, B. A. Reynolds and L. P. Deleyrolle (2018). “Infiltrative and drug-resistant slow-cycling cells support metabolic heterogeneity in glioblastoma.” EMBO J.

Hochgerner, H., A. Zeisel, P. Lonnerberg and S. Linnarsson (2018). “Conserved properties of dentate gyrus neurogenesis across postnatal development revealed by single-cell RNA sequencing.” Nat Neurosci 21(2): 290–299.

Jahn, H. M., C. V. Kasakow, A. Helfer, J. Michely, A. Verkhratsky, H. H. Maurer, A. Scheller and F. Kirchhoff (2018). “Refined protocols of tamoxifen injection for inducible DNA recombination in mouse astroglia.” Sci Rep 8(1): 5913.

Jimenez-Pascual, A., J. S. Hale, A. Kordowski, J. Pugh, D. J. Silver, D. Bayik, G. Roversi, T. J. Alban, S. Rao, R. Chen, T. M. McIntyre, G. Colombo, G. Taraboletti, K. O. Holmberg, K. Forsberg-Nilsson, J. D. Lathia and F. A. Siebzehnrubl (2019). “ADAMDEC1 Maintains a Growth Factor Signaling Loop in Cancer Stem Cells.” Cancer Discov 9(11): 1574–1589.

Kempermann, G., F. H. Gage, L. Aigner, H. Song, M. A. Curtis, S. Thuret, H. G. Kuhn, S. Jessberger, P. W. Frankland, H. A. Cameron, E. Gould, R. Hen, D. N. Abrous, N. Toni, A. F. Schinder, X. Zhao, P. J. Lucassen and J. Frisen (2018). “Human Adult Neurogenesis: Evidence and Remaining Questions.” Cell Stem Cell 23(1): 25–30.

Kempermann, G., S. Jessberger, B. Steiner and G. Kronenberg (2004). “Milestones of neuronal development in the adult hippocampus.” Trends Neurosci 27(8): 447–452.

Knoblich, J. A. (2008). “Mechanisms of asymmetric stem cell division.” Cell 132(4): 583–597.

Kuwabara, T., J. Hsieh, A. Muotri, G. Yeo, M. Warashina, D. C. Lie, L. Moore, K. Nakashima, M. Asashima and F. H. Gage (2009). “Wnt-mediated activation of NeuroD1 and retro-elements during adult neurogenesis.” Nat Neurosci 12(9): 1097–1105.

Lazutkin, A., O. Podgorny and G. Enikolopov (2019). “Modes of division and differentiation of neural stem cells.” Behav Brain Res 374: 112118.

Ligon, K. L., E. Huillard, S. Mehta, S. Kesari, H. Liu, J. A. Alberta, R. M. Bachoo, M. Kane, D. N. Louis, R. A. Depinho, D. J. Anderson, C. D. Stiles and D. H. Rowitch (2007). “Olig2-regulated lineage-restricted pathway controls replication competence in neural stem cells and malignant glioma.” Neuron 53(4): 503–517.

Liu, J., Y. Liu, J. Shao, Y. Li, L. Qin, H. Shen, Y. Xie, W. Xia and W. Q. Gao (2019). “Zeb1 is important for proper cleavage plane orientation of dividing progenitors and neuronal migration in the mouse neocortex.” Cell Death Differ 26(11): 2479–2492.

Liu, Y., L. Siles, X. Lu, K. C. Dean, M. Cuatrecasas, A. Postigo and D. C. Dean (2018). “Mitotic polarization of transcription factors during asymmetric division establishes fate of forming cancer cells.” Nat Commun 9(1): 2424.

Madisen, L., T. A. Zwingman, S. M. Sunkin, S. W. Oh, H. A. Zariwala, H. Gu, L. L. Ng, R. D. Palmiter, M. J. Hawrylycz, A. R. Jones, E. S. Lein and H. Zeng (2010). “A robust and high-throughput Cre reporting and characterization system for the whole mouse brain.” Nat Neurosci 13(1): 133–140.

Maekawa, M., N. Takashima, Y. Arai, T. Nomura, K. Inokuchi, S. Yuasa and N. Osumi (2005). “Pax6 is required for production and maintenance of progenitor cells in postnatal hippocampal neurogenesis.” Genes Cells 10(10): 1001–1014.

Martin-Suarez, S., J. Valero, T. Muro-Garcia and J. M. Encinas (2019). “Phenotypical and functional heterogeneity of neural stem cells in the aged hippocampus.” Aging Cell 18(4): e12958.

Miller, I., M. Min, C. Yang, C. Tian, S. Gookin, D. Carter and S. L. Spencer (2018). “Ki67 is a Graded Rather than a Binary Marker of Proliferation versus Quiescence.” Cell Rep 24(5): 1105–1112 e1105.

Ming, G. L. and H. Song (2011). “Adult neurogenesis in the mammalian brain: significant answers and significant questions.” Neuron 70(4): 687–702.

Mori, T., K. Tanaka, A. Buffo, W. Wurst, R. Kuhn and M. Gotz (2006). “Inducible gene deletion in astroglia and radial glia--a valuable tool for functional and lineage analysis.” Glia 54(1): 21–34.

Palmer, T. D., J. Takahashi and F. H. Gage (1997). “The adult rat hippocampus contains primordial neural stem cells.” Mol Cell Neurosci 8(6): 389–404.

Peinado, H., D. Olmeda and A. Cano (2007). “Snail, Zeb and bHLH factors in tumour progression: an alliance against the epithelial phenotype?” Nat Rev Cancer 7(6): 415–428.

Pilz, G. A., S. Bottes, M. Betizeau, D. J. Jorg, S. Carta, B. D. Simons, F. Helmchen and S. Jessberger (2018). “Live imaging of neurogenesis in the adult mouse hippocampus.” Science 359(6376): 658–662.

Rosmaninho, P., S. Mukusch, V. Piscopo, V. Teixeira, A. A. Raposo, R. Warta, R. Bennewitz, Y. Tang, C. Herold-Mende, S. Stifani, S. Momma and D. S. Castro (2018). “Zeb1 potentiates genome-wide gene transcription with Lef1 to promote glioblastoma cell invasion.” EMBO J 37(15).

Ryu, J. R., C. J. Hong, J. Y. Kim, E. K. Kim, W. Sun and S. W. Yu (2016). “Control of adult neurogenesis by programmed cell death in the mammalian brain.” Mol Brain 9: 43.

Seri, B., J. M. Garcia-Verdugo, B. S. McEwen and A. Alvarez-Buylla (2001). “Astrocytes give rise to new neurons in the adult mammalian hippocampus.” J Neurosci 21(18): 7153–7160.

Siebzehnrubl, F. A., D. J. Silver, B. Tugertimur, L. P. Deleyrolle, D. Siebzehnrubl, M. R. Sarkisian, K. G. Devers, A. T. Yachnis, M. D. Kupper, D. Neal, N. H. Nabilsi, M. P. Kladde, O. Suslov, S. Brabletz, T. Brabletz, B. A. Reynolds and D. A. Steindler (2013). “The ZEB1 pathway links glioblastoma initiation, invasion and chemoresistance.” EMBO Mol Med 5(8): 1196–1212.

Sierra, A., J. M. Encinas, J. J. Deudero, J. H. Chancey, G. Enikolopov, L. S. Overstreet-Wadiche, S. E. Tsirka and M. Maletic-Savatic (2010). “Microglia shape adult hippocampal neurogenesis through apoptosis-coupled phagocytosis.” Cell Stem Cell 7(4): 483–495.

Singh, D. K., R. K. Kollipara, V. Vemireddy, X. L. Yang, Y. Sun, N. Regmi, S. Klingler, K. J. Hatanpaa, J. Raisanen, S. K. Cho, S. Sirasanagandla, S. Nannepaga, S. Piccirillo, T. Mashimo, S. Wang, C. G. Humphries, B. Mickey, E. A. Maher, H. Zheng, R. S. Kim, R. Kittler and R. M. Bachoo (2017). “Oncogenes Activate an Autonomous Transcriptional Regulatory Circuit That Drives Glioblastoma.” Cell Rep 18(4): 961–976.

Singh, S., D. Howell, N. Trivedi, K. Kessler, T. Ong, P. Rosmaninho, A. A. Raposo, G. Robinson, M. F. Roussel, D. S. Castro and D. J. Solecki (2016). “Zeb1 controls neuron differentiation and germinal zone exit by a mesenchymal-epithelial-like transition.” Elife 5.

Spaderna, S., O. Schmalhofer, M. Wahlbuhl, A. Dimmler, K. Bauer, A. Sultan, F. Hlubek, A. Jung, D. Strand, A. Eger, T. Kirchner, J. Behrens and T. Brabletz (2008). “The transcriptional repressor ZEB1 promotes metastasis and loss of cell polarity in cancer.” Cancer research 68(2): 537–544.

Steiner, B., G. Kronenberg, S. Jessberger, M. D. Brandt, K. Reuter and G. Kempermann (2004). “Differential regulation of gliogenesis in the context of adult hippocampal neurogenesis in mice.” Glia 46(1): 41–52.

Stemmler, M. P., R. L. Eccles, S. Brabletz and T. Brabletz (2019). “Non-redundant functions of EMT transcription factors.” Nat Cell Biol 21(1): 102–112.

Suh, H., A. Consiglio, J. Ray, T. Sawai, K. A. D’Amour and F. H. Gage (2007). “In vivo fate analysis reveals the multipotent and self-renewal capacities of Sox2+ neural stem cells in the adult hippocampus.” Cell Stem Cell 1(5): 515–528.

Takagi, T., H. Moribe, H. Kondoh and Y. Higashi (1998). “DeltaEF1, a zinc finger and homeodomain transcription factor, is required for skeleton patterning in multiple lineages.” Development 125(1): 21–31.

Trent, S., J. Hall, W. M. Connelly and A. C. Errington (2019). “Cyfip1 Haploinsufficiency Does Not Alter GABAA Receptor delta-Subunit Expression and Tonic Inhibition in Dentate Gyrus PV(+) Interneurons and Granule Cells.” eNeuro 6(3).

Wang, H., Z. Xiao, J. Zheng, J. Wu, X. L. Hu, X. Yang and Q. Shen (2019). “ZEB1 Represses Neural Differentiation and Cooperates with CTBP2 to Dynamically Regulate Cell Migration during Neocortex Development.” Cell Rep 27(8): 2335–2353 e2336.

White, C. W., 3rd, X. Fan, J. C. Maynard, E. G. Wheatley, G. Bieri, J. Couthouis, A. L. Burlingame and S. A. Villeda (2020). “Age-related loss of neural stem cell O-GlcNAc promotes a glial fate switch through STAT3 activation.” Proc Natl Acad Sci U S A 117(36): 22214–22224.

Ziebell, F., A. Martin-Villalba and A. Marciniak-Czochra (2014). “Mathematical modelling of adult hippocampal neurogenesis: effects of altered stem cell dynamics on cell counts and bromodeoxyuridine-labelled cells.” J R Soc Interface 11(94): 20140144.

